# Genome-wide Profiling of Histone Modifications in *Plasmodium falciparum* using CUT&RUN

**DOI:** 10.1101/2022.08.01.502378

**Authors:** Riward Campelo Morillo, Chantal T. Harris, Kit Kennedy, Björn F.C. Kafsack

**Affiliations:** Department of Microbiology & Immunology, Weill Cornell Medicine, New York, NY, USA; Immunology and Microbial Pathogenesis Program

## Abstract

We recently adapted a CUT&RUN protocol for genome-wide profiling of chromatin modifications in the human malaria parasite *Plasmodium*. Using the step-by-step protocol described below, we were able to generate high quality profiles of multiple histone modifications using only a small fraction of the cells required for ChIPseq. Using antibodies against two commonly profiled histone modifications, H3K4me3 and H3K9me3, we show here that CUT&RUN profiling is highly reproducible and closely recapitulates previously published ChIPseq-based abundance profiles of histone marks. Finally, we show that CUT&RUN requires substantially lower sequencing coverage for accurate profiling compared to ChIPseq.

## Introduction

Post-translational histone modifications play a central role in regulating chromatin accessibility and partitioning in nearly all eukaryotes. Numerous modifications have been described with a wide variety of functions and dynamics (reviewed in Rothbart & Strahl 2014). The ability to profile these modifications genome-wide is a key tool for assigning their role in regulating gene expression and, more generally, for investigating nuclear functions. Traditionally this has been accomplished by Chromatin ļmmunoPrecipitation (ChIP) (Solomon et al. 1988), which relies on affinity-based pull-down from randomly fragmented chromatin of protein-DNA complexes with antibodies specific to one of the complex components followed by the extraction of the genomic fragments. High Throughput Sequencing (HTS) methods are then used to determine the relative population abundance of the antibody target at each genomic position based on its relative enrichment in the precipitated chromatin versus its genomic frequency.

ChIP-seq has several drawbacks that derive from the random fragmentation of chromatin and the requirement for stringency during immunoprecipitation. The former results in relatively high background while both make profiling of low affinity interactions difficult. This can be overcome by protein-DNA cross-linking prior to fragmentation. However, crosslinking presents another challenge as it can lead to non-specific recovery of untargeted proteins correlated with duration of fixation treatment (Baranello et al. 2016) and interfere with antibody recognition by epitope disruption (O’Neill & Turner 2003).

The recent development of <u>C</u>leavage <u>U</u>nder <u>T</u>argets & <u>R</u>elease <u>U</u>sing <u>N</u>uclease (CUT&RUN) addresses many of these challenges (Figure 1) (Skene et al. 2018; Meers et al. 2019). Instead of isolating and randomly fragmenting chromatin, cells are immobilized on beads and permeabilized to allow antibodies to diffuse into the nucleus to bind their targets in situ. After removal of excess antibodies, a fusion of protein A/G, which binds the Fc portion of antibodies, and micrococcal nuclease (MNase) is added under low calcium conditions that keep the MNase inactive. Following the removal of excess proteinA/G-MNase, calcium is added to activate MNase cleavage thereby releasing the antibody-bound nucleosomes but leaving the unbound fraction of the genome intact. Due to their small size, liberated nucleosomes can then diffuse out of the cell while the rest of the genome remains cell-associated. DNA can then be extracted from the supernatant for standard HTS library generation, high-throughput sequencing, and analysis. Because of this CUT&RUN has exceptionally low background and has been successfully used to profile chromatin interactions from very small numbers of cells (Skene et al. 2018; Liu et al. 2018). CUT&RUN has been successfully used for profiling a wide-array of chromatin proteins in many experimental systems. In this study we specifically focus on its use for profiling post-translational histone modifications on nucleosome.

**Figure 1.**
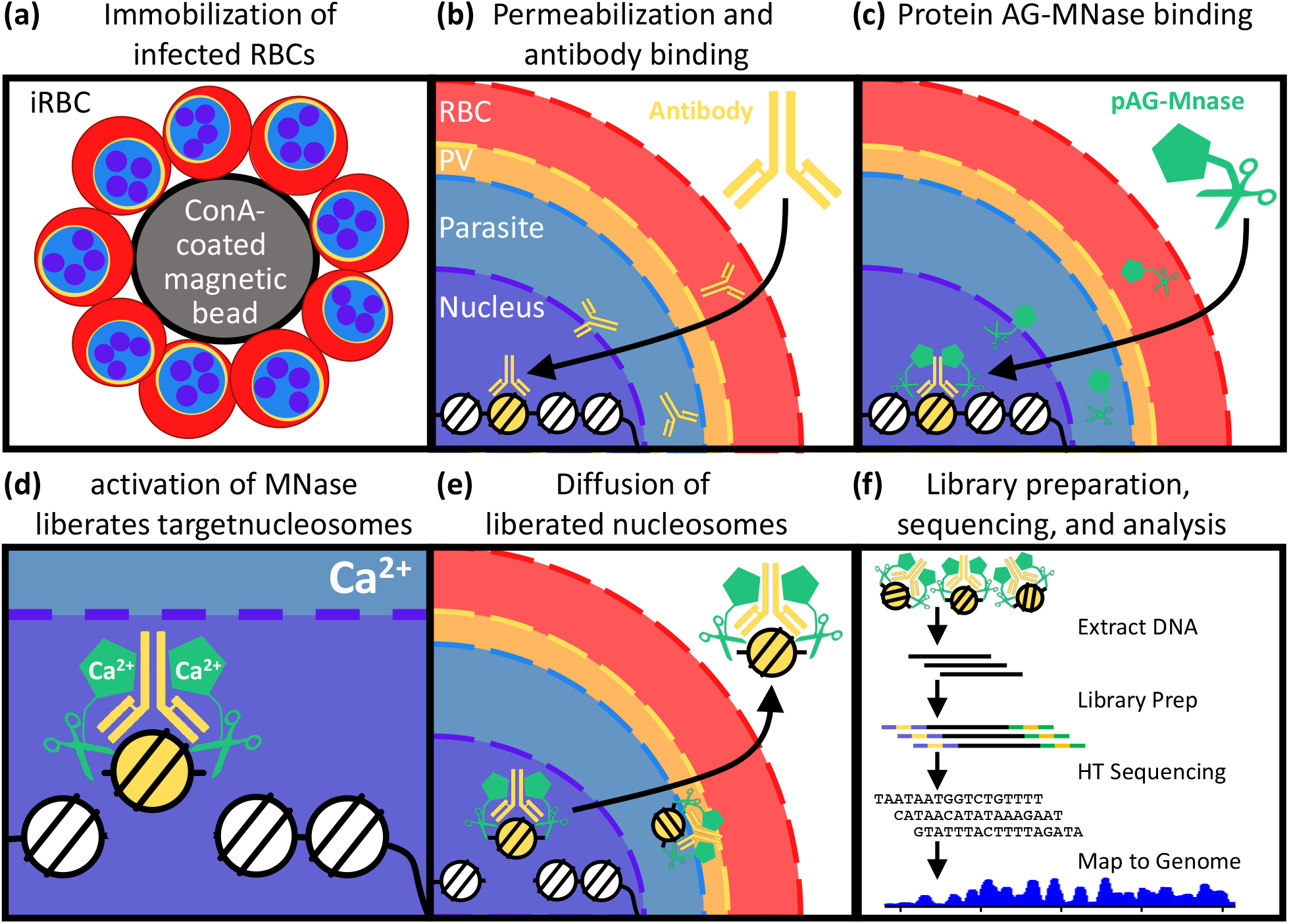
Overview of genome-wide profiling of histone modifications using CUT & RUN. **(a)** Infected RBCs are immobilized on Concavalin-coated beads. **(b)** Immobilized cells are permeabilized allowing histone modification-specific antibody to diffuse into the nucleus to bind its targets. **(c)** After removal of excess antibodies, MNase fused to proteinAG is added under low Calcium conditions and binds to chromatin-associated antibodies. **(d)** After removal of excess proteinAG-MNase, cleavage adjacent to bound nucleosomes is activated by addition of Calcium cations. **(e)** liberated nucleosomes are **a**llowed to diffuse out of the permeabilized cells for subsequent **(f)** DNA extraction, library generation, high-throughput sequencing, and analysis.

We recently adapted a CUT&RUN protocol (Meers et al. 2019, dx.doi.org/10.17504/protocols.io.zcpf2vn) for genome-wide profiling of chromatin modifications in the human malaria parasite *Plasmodium falciparum* (Harris et al. 2022). Using the step-by-step protocol described below, we were able to generate high quality profiles of multiple histone modifications using only a small fraction of the cells required for ChIPseq. Using antibodies against two commonly profiled histone modifications, H3K4me3 and H3K9me3, we show here that CUT&RUN profiling is highly reproducible (Figure 2, Supplementary Figure 1) and closely recapitulates previously published ChIPseq-based abundance profiles of histone marks (Figure 3, Supplementary Figures 1-2) (Bunnik et al. 2018; Fraschka et al. 2018; Bartfai et al. 2010). Finally, we show that CUT&RUN requires substantially lower sequencing coverage for accurate profiling compared to ChIP-seq (Figure 4).

**Figure 2.**
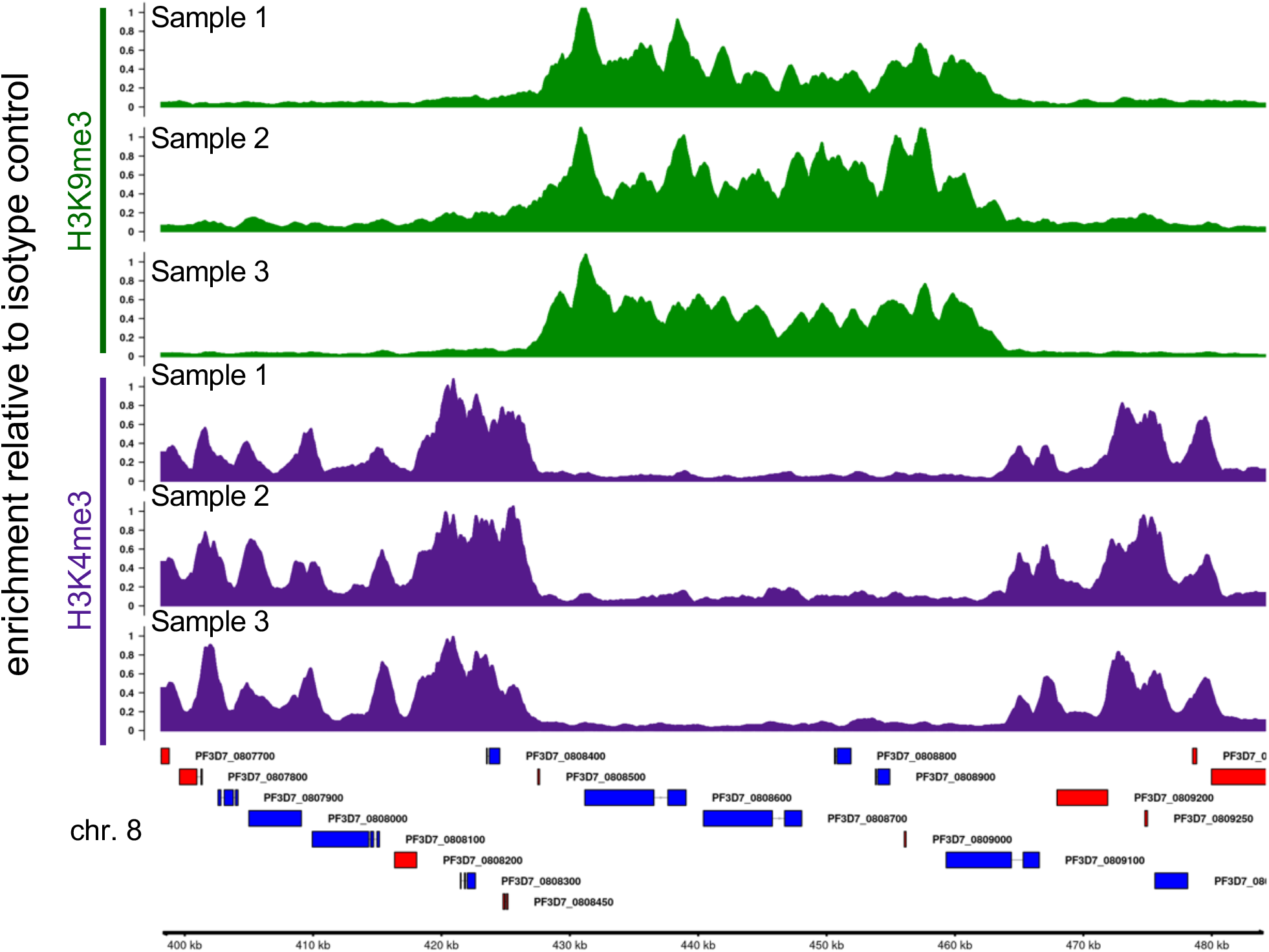
CUT&RUN profiles of histone modifications are highly reproducible. Fold Enrichment profiles of H3K9me3 (green) and H3K4me3 (purple) in three biological replicates. H3K4me3 and H3K9me3 tracks were scaled to maximal value in the region shown. Example locus on chromosome 8 contains a non-subtelomeric heterochromatin island. Genes shown in blue are encoded on the top (+) strand while those in red are encoded on the bottom (-) strand. Gene IDs are shown to the right of the gene.

**Figure 3.**
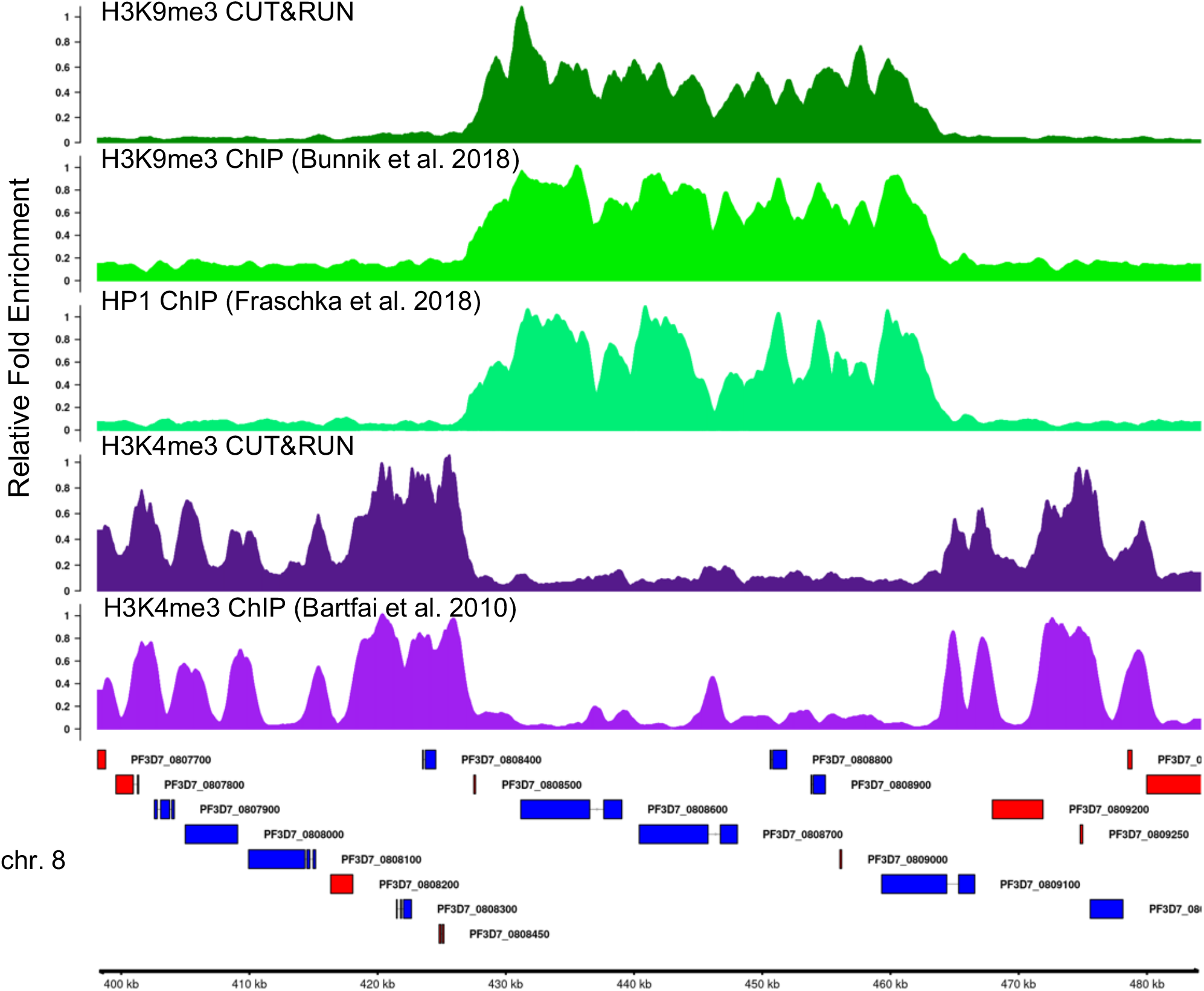
Comparison of H3K4me3 and H3K9me3 profiles obtained by CUT&RUN to published ChIP-seq profiles at an example locus containing a non-subtelomeric heterochromatin island on chromosome 8. Tracks show the relative fold-enrichment of specific histone modifications, H3K9me3 & H3K4me3, as well as for heterochromatin protein 1, the H3K9me3 histone reader, versus either isotype controls for CUT&RUN or input DNA (ChIPseq). Sequence reads from published stage-matched H3K9me3 (SRR4444647, SRR4444639), HP1 (SRR5935737, SRR5935738), and H3K4me3 (SRR065659, SRR065664) ChIP-seq experiments were downloaded from the NCBI Sequence Read Archive. Genes shown in blue are encoded on the top (+) strand while those in red are encoded on the bottom (-) strand. GeneIDs are shown to the right of each gene.

**Figure 4.**
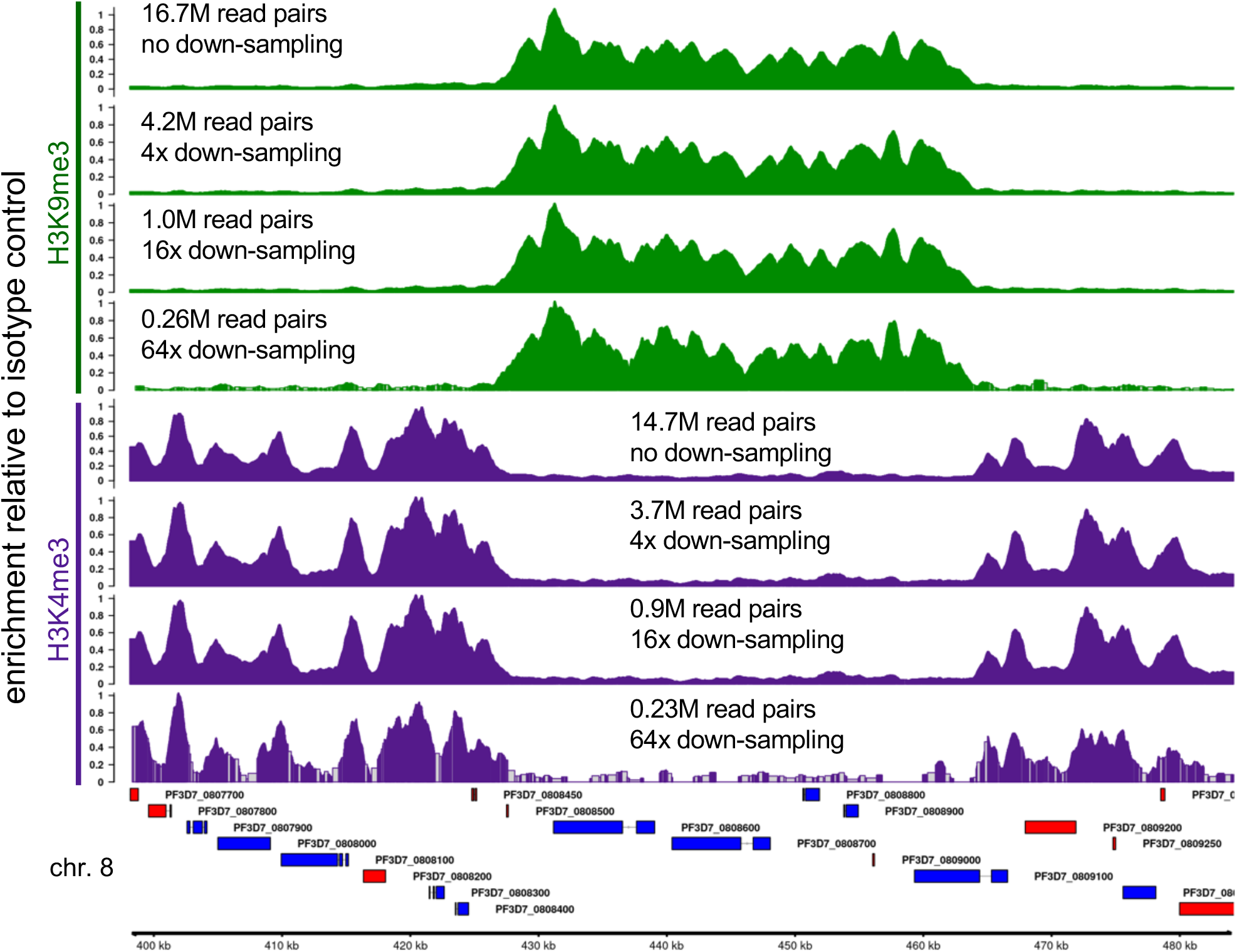
Fold Enrichment profiles of down-sampled H3K9me3 (green) and H3K4me3 (purple) CUT&RUN reads show good signal-to-noise ratios down to ^~^1 million read-pairs per sample. Read pairs were serially down-sampled *in silico* by 4-fold to evaluate the signal-to-noise ratio. Each track is scaled to its maximal value across the locus shown. The number of sequenced paired-end 50bp (PE50) clusters used are indicated in millions. Example locus on chromosome 8 contains a non-subtelomeric heterochromatin island. Genes shown in blue are encoded on the top (+) strand while those in red are encoded on the bottom (-) strand. GeneIDs are shown the above or below the gene. Grey regions in the 64× down-sampled tracks lack coverage and are inferred based on neighboring bins.

## Planning and Preparation

### Prepare stock solutions

**Timing: 4 hours**

**1 M HEPES-NaOH pH 7.5:** Dissolve 238.3 g HEPES in 800 mL of distilled water. Adjust pH to 7.5 using 1M NaOH then adjust the final volume with distilled water to 1 L. Store at RT.

**1 M HEPES-KOH:** Dissolve 238. 3 g HEPES in 800 mL of distilled water. Adjust pH to 7.5 using 1M KOH then adjust the final volume with distilled water to 1 L. Store at RT.

**1 M KCl:** Dissolve 74.55 g KCl in 800 mL of distilled water and adjust the volume to 1 L. Store at RT.

**5% Sorbitol**: Dissolve 50 g Sorbitol in 800 mL of distilled water and adjust the volume to 1 L.

Filter sterilize and store at RT.

**90% Percoll with 6% Sorbitol:** Mix 20 mL minimal RPMI with 180 mL Percoll and dissolve 12 g D-sorbitol. Filter sterilize and store at 4°C.

**100 mM CaCl2:** Dissolve 1.1 g CaCl2 in 100 mL of distilled water.

**100 mM MnCl2:** Dissolve 1.25 g MnCl2 in 100 mL of distilled water.

**100 mM EGTA pH 8.0:** Dissolve 3.8 g EGTA in 80 mL of distilled water while maintaining the pH at 8.0 using 10N NaOH. Adjust volume to 100 mL and store at RT.

**500 mM EDTA pH 8.0:** Dissolve 186.1 g EDTA in 800 mL distilled water while maintaining the pH at 8.0 using 10N NaOH. Adjust volume to 1 L and store at RT.

**1 M Tris-HCL pH 8.0:** Dissolve 121.14 g Tris in 800 mL of distilled water Adjust pH to 8.0 using concentrated HCl / 10N NaOH. Adjust the final volume with distilled water to 1 L and store at RT.

**5 M NaCl:** Dissolve 292.2 g NaCl in 800 mL distilled water. Adjust the volume to 1 L and store at RT.

**5% Digitonin:** Dissolve 0.05 g of digitonin in 1 mL of dimethylsulfoxide (DMSO). Prepare 150 μL aliquots and store at −20 °C for up to 6 months.

**25× Protease Inhibitor cocktail:** Dissolve 1 EDTA-free cOmplete protease inhibitor tablet in 1 mL of distilled water. Split into 200μL aliquots and store at −20 °C for up to 3 months.

**2 M spermidine:** Prepare a *fresh* work solution each time by diluting 31.25 μL of stock spermidine (6.4 M) in 100 μl of distilled water.

### *P. falciparum* culturing and synchronization of erythrocytic stages

**Timing: ^~^1 week**

We have found that CUT&RUN using parasites containing 10-50 million nuclei yielded robust signal for the antibodies tested. At a parasitemia and hematocrit of 5% each, this is the equivalent of 0.4-2mL of culture for 1N parasites (ring, early trophozoite, and gametocytes stages) or as little as 20-100 μL of culture containing segmenting schizonts.

1. Maintain parasite cultures in T25 flasks following established culturing techniques (Moll et al. 2008) using red blood cells at 3-5% hematocrit resuspended in standard malaria complete media (MCM).
2. Double-synchronize parasite cultures with sorbitol treatment to achieve a synchrony of ± 6 h (Moll et al. 2008).
  a. Wash away media by centrifugation 800*g* × 3 min at RT.
  b. Resuspend pelleted RBC by gently adding 10× volumes of 5% sorbitol.
  c. Incubate for 5-10 min in a water bath at 37°C.
  d. Spin down to wash away sorbitol and resuspend in fresh media.

## Reagents, Materials, and Equipment

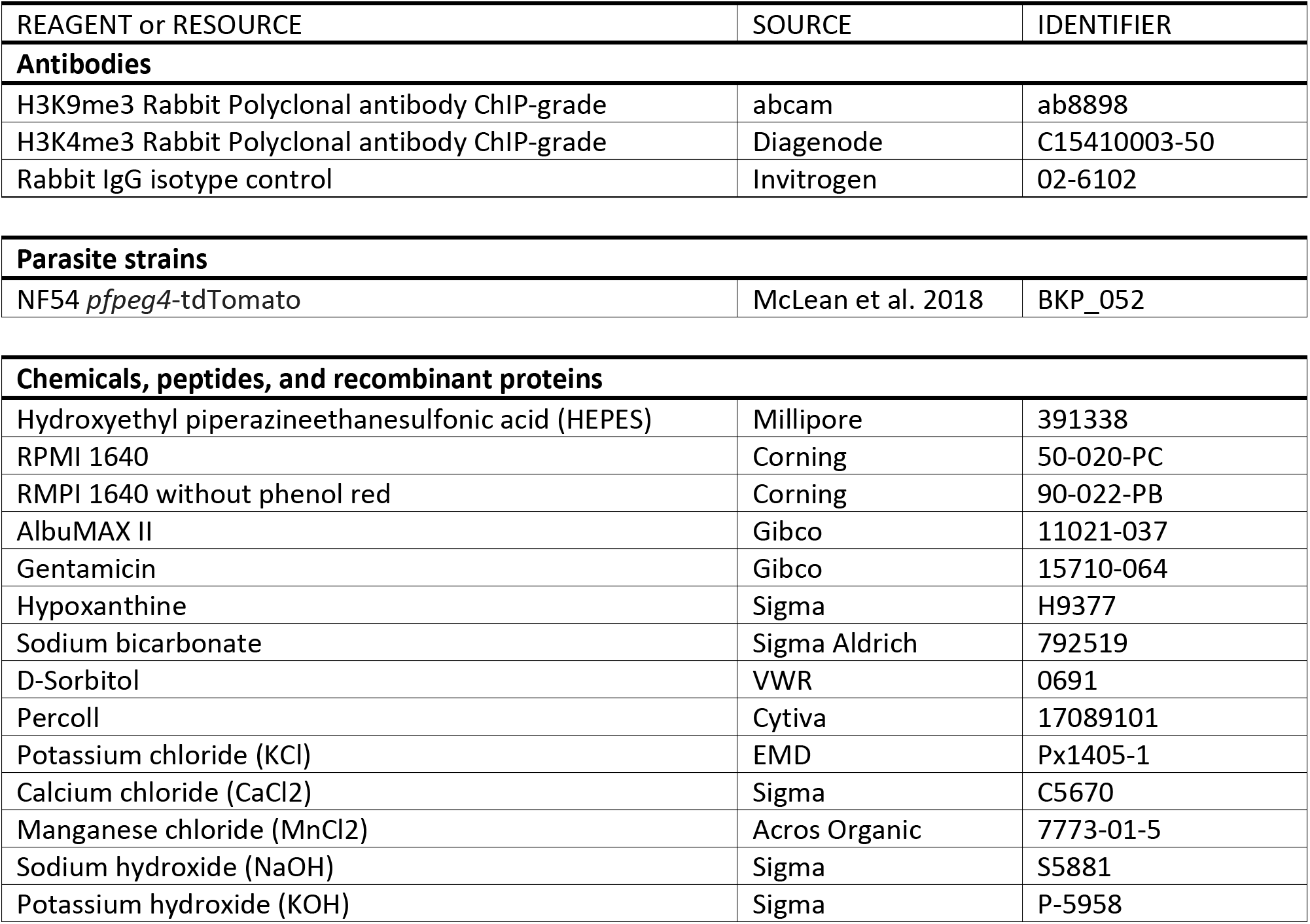
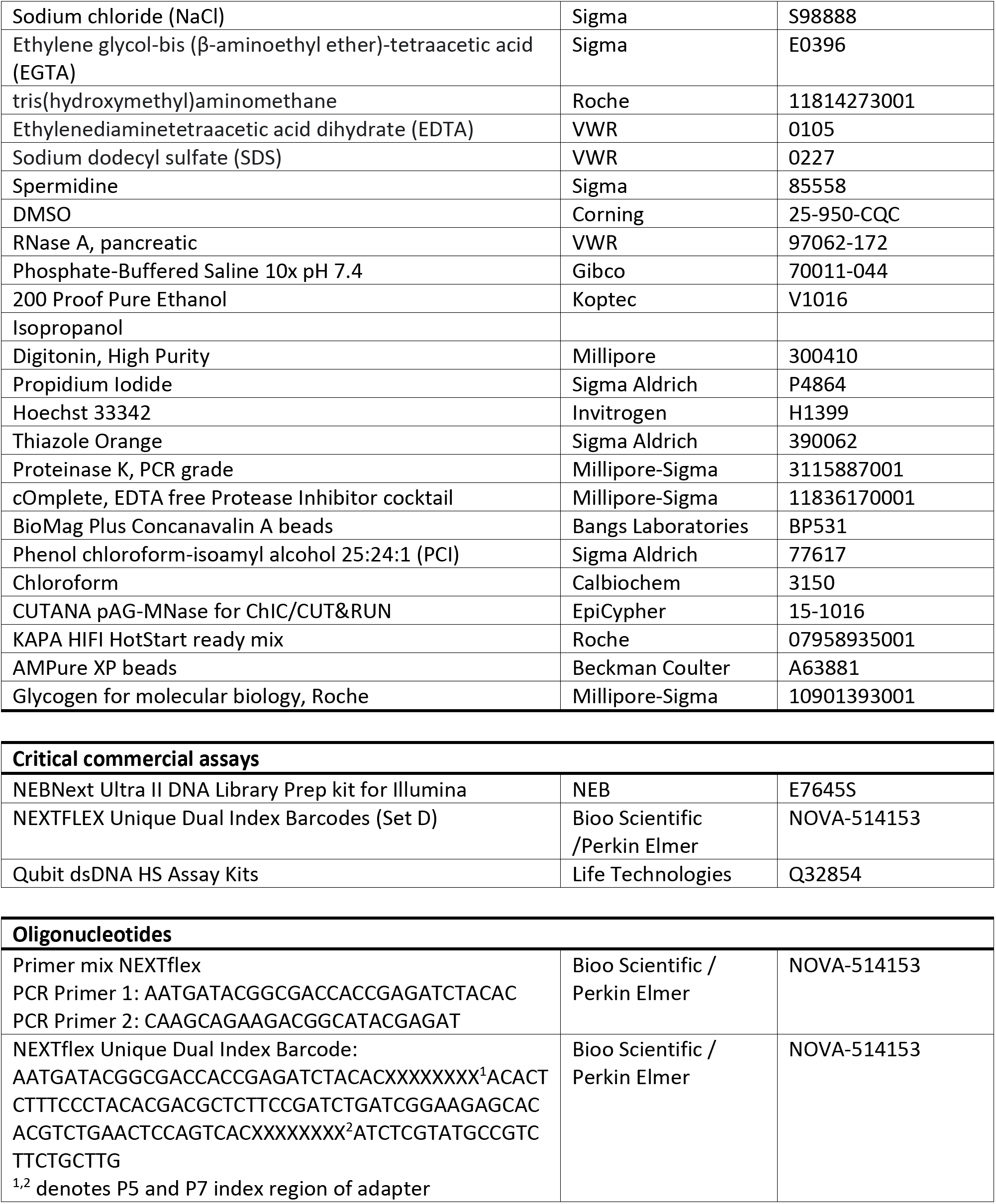
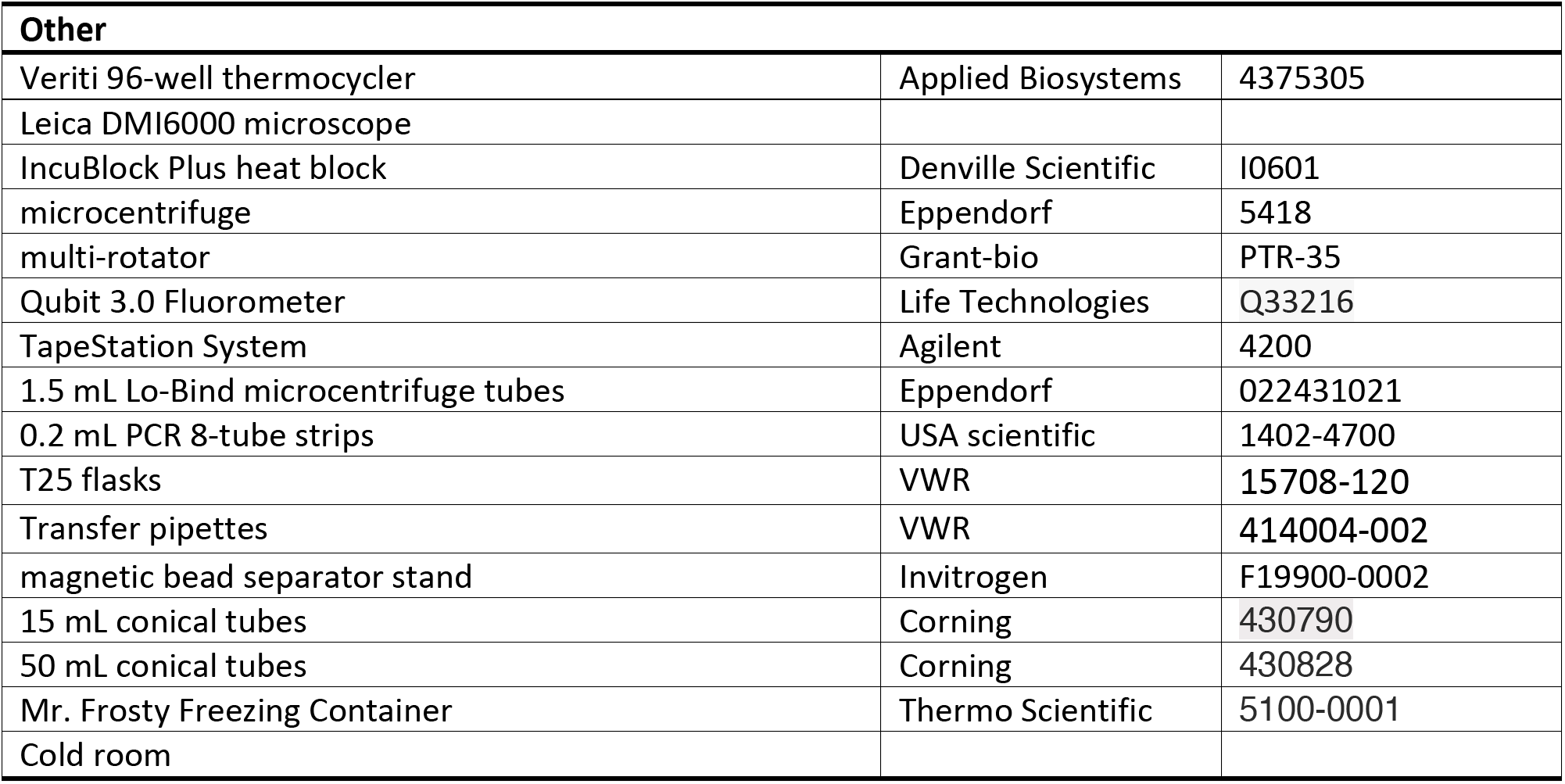

### Solutions and Buffers

#### Binding buffer

Solutions and Buffers **Binding buffer**
|  | **Final concentration** | **Amount** |
| --- | --- | --- |
| HEPES-KOH pH 7.5 (1 M) | 20 mM | 400 µL |
| KCl (1 M) | 10 mM | 200 µL |
| CaCl<sub>2</sub> (100 mM) | 1 mM | 200 µL |
| MnCl<sub>2</sub> (100 mM) | 1 mM | 200 µL |
| ddH<sub>2</sub>O | n/a | 19 mL |
| **Total** | **n/a** | **20 mL** |

Store at 4°C for up to 6 months. *Note that the Binding buffer uses HEPES-KOH!*

#### Wash buffer

|  | **Final concentration** | **Amount** |
| --- | --- | --- |
| HEPES-NaOH pH 7.5 (1 M) | 20 mM | 1 mL |
| NaCl (5 M) | 150 mM | 1.5 mL |
| Spermidine (2 M) | 0.5 mM | 12.5 µL |
| ddH<sub>2</sub>O | n/a | 45.5 mL |
| **Total** | **n/a** | **50 mL** |

Store at 4°C for up to 1 week

**CRITICAL:** *Protease inhibitor cocktail (EDTA-free) should be added fresh on the day of use*.Add 20 μl of Protease inhibitor cocktail stock solution per 1 mL of **Wash buffer**.

#### Dig-Wash buffer

|  | Final concentration | Amount |
| --- | --- | --- |
| **Wash buffer** | n/a | 30 mL |
| Digitonin (5%) | 0.025 % (See Note) | 150 μL |
| **Total** | **n/a** | **30 mL** |

Store on ice or at 4°C for up to 1 day

**CRITICAL:** *Mix well before each use*.

**Note:** Digitonin has batch variability and the minimal concentration that yields full of permeabilization needs to be determined empirically for your digitonin stock. We have obtained good results with as low as 0.025% digitonin. To test for permeabilization, pellet 150 μL of culture, and resuspend in 150μL **Dig-Wash buffer** containing 8 μM Hoechst 33342 and 10 μg/mL propidium iodide. Incubate cell suspension for 10 min at RT and wash once. Pellet and resuspend cells in 20 μL of media and observe under fluorescence microscope or flow cytometer. Determine the lowest digitonin concentration at which all nuclei are positive for both dyes.

**Figure 5.**
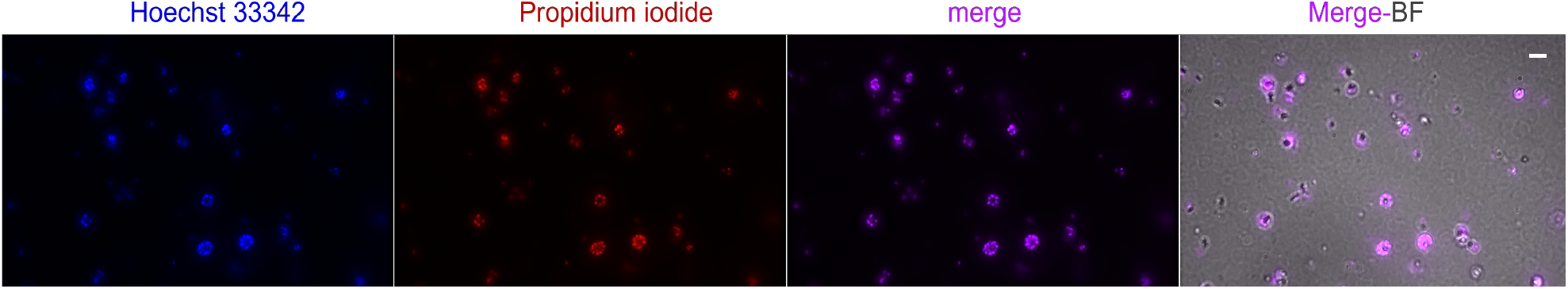
iRBCs treated with 0.025 % Digitonin and stained with Hoechst 33342 and propidium iodide. Nuclei of permeabilized cells will stain with both dyes. Scale bar, 4 μm.

#### Antibody buffer

|  | Final concentration | Amount |
| --- | --- | --- |
| **Dig-Wash buffer** | n/a | 2 mL |
| EDTA pH 8.0 (0.5 M) | 2 mM | 8 μL |
| **Total** | **n/a** | **2 mL** |

*Always prepare fresh before use*.

**CRITICAL:** *Add the amount of antibody indicated right before incubation and keep on ice at all times*.

#### Low-Salt Rinse buffer

|  | Final concentration | Amount |
| --- | --- | --- |
| HEPES-NaOH pH 7.5 (1 M) | 20 mM | 400 μL |
| Digitonin (5%) | 0.025 % | 100 μL |
| Spermidine (2 M) | 0.5 mM | 12.5 μL |
| ddH<sub>2</sub>O | n/a | 19.5 mL |
| **Total** | **n/a** | **20 mL** |

Store at 4°C for up to 1 week.

**CRITICAL:** *Protease inhibitor cocktail (EDTA-free) should be added fresh on the day of use*.Add 20 μl of 25× stock solution per 1 mL of buffer.

#### High Ca^++^ Incubation buffer

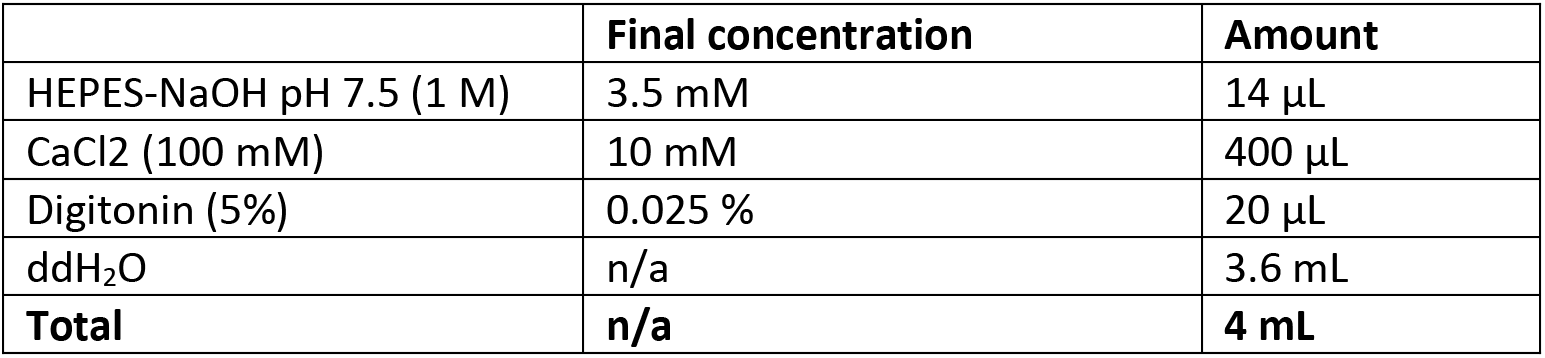

|  | Final concentration | Amount |
| --- | --- | --- |
| HEPES-NaOH pH 7.5 (1 M) | 3.5 mM | 14 µL |
| CaCl<sub>2</sub> (100 mM) | 10 mM | 400 µL |
| Digitonin (5%) | 0.025 % | 20 µL |
| ddH<sub>2</sub>O | n/a | 3.6 mL |
| **Total** | **n/a** | **4 mL** |

Store at 4°C for up to 1 week.

**CRITICAL:** *Protease inhibitor cocktail (EDTA-free) should be added fresh on the day of use*.

Add 20 μl of 25× stock solution per 1 mL of buffer.

#### Stop buffer

|  | Final concentration | Amount |
| --- | --- | --- |
| NaCl (5 M) | 170 mM | 170 µL |
| EGTA (100 mM) | 20 mM | 1 mL |
| Digitonin (5%) | 0.025 % | 25 µL |
| RNase A (10 mg/mL) | 50 µg/mL | 12.5 µL |
| Glycogen (20 mg/mL) | 25 µg/mL | 6.25 µL |
| ddH<sub>2</sub>O | n/a | 3.8 mL |
| **Total** | **n/a** | **5 mL** |

Store at 4°C for up to 1 week.

#### TE buffer

|  | Final concentration | Amount |
| --- | --- | --- |
| Tris-HCl (1 M) | 10 mM | 1 mL |
| EDTA (500 mM) | 1 mM | 0.2 mL |
| ddH<sub>2</sub>O | n/a | 98.8 mL |
| **Total** | **n/a** | **100 mL** |

### Step-by-step Instructions

#### Activation of concanavalin A-coated beads

**Timing: 5 minutes**

*Concanavalin A lectins on magnetic beads are activated and stabilized by Ca^2+^ and Mn^2+^ cations*.

1. Gently resuspend and withdraw 28 μL of the ConA bead slurry per sample/antibody combination.
2. Transfer beads to a 1.5 mL microcentrifuge tube and add 1 mL of **Binding buffer**.
3. Place the tube on a magnetic bead separator stand and allow to clear (30-120 sec) before removing and discarding the supernatant.
4. Remove tube from stand and add 1 mL of **Binding buffer**. Mix gently and return to the magnetic stand.
5. Repeat steps 3-4 one additional time.
6. Resuspend beads in the same volume of **Binding Buffer** that was added in step 1 and keep at RT.
7. Place **Wash Buffer** at RT to warm for later use.

#### Binding cells to Concanavalin A-coated beads

**Timing: 1 hour**

*Infected red blood cells (iRBCs) bind to beads coated with concanavalin A, a lectin that binds specifically to extracellular glycoproteins*.

8. Harvest fresh parasite cultures at the desired stage and synchrony. If feasible, enrich for iRBCs by magnetic purification (Mata-Cantero et al. 2014), centrifugation in a continuous percoll gradient (Rivadeneira et al. 1983), or lysis of uninfected RBCs (Brown et al. 2020). The stage, synchrony and growth conditions are determined by the scientific question the experimenter is asking. In this study, we used percoll/sorbitol isolated asexual blood stages synchronized to a window of 36±4 hours post-invasion.
9. Determine iRBC density and purity using a hemocytometer, coulter counter, or volumetric flow cytometry.
10. Pellet cells by centrifugation (800*g* for 3 min at RT) and resuspend cells containing 1-5×10^7^ nuclei to a cell density to 1×10^7^ cells/mL in **Wash buffer** by gentle pipetting.

**Note:** The average nuclear content of parasites can be determined by flow cytometry after staining cultures with 8 μM Hoechst 33342 (DNA-specific) and 0.1 μg/mL thiazole orange (RNA-specific with Hoechst 33342) for 30 min at 37 ° C, followed by two brief washes. A high ring-stage culture can be used to determine the mean fluorescence value equivalent to a 1N nuclear content. However, we did not attempt to optimize the minimal number of nuclei required. Based on CUT&RUN in other systems and our down-sampling analysis (Figure 4) substantially fewer nuclei are likely sufficient.

**Figure 6.**
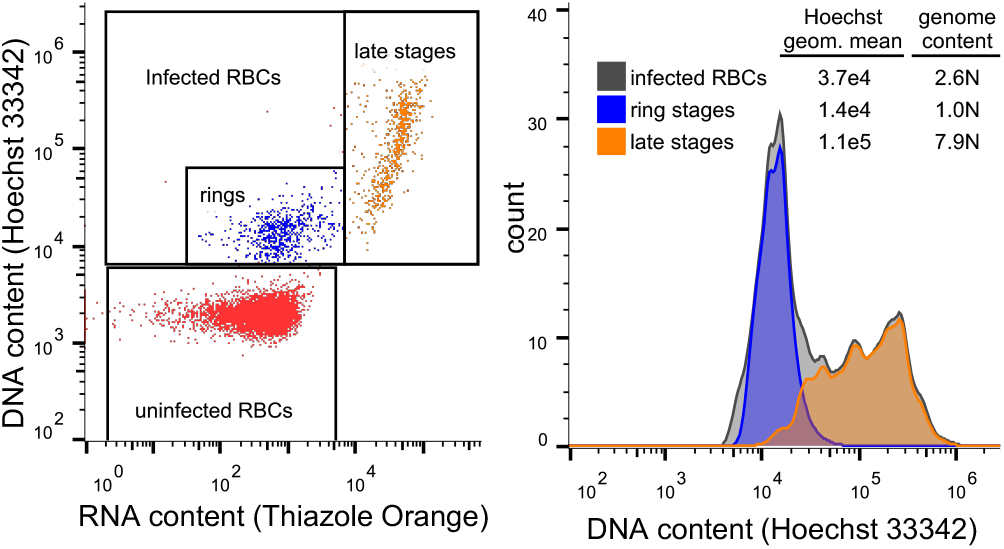
Quantification of parasite genome content. Left: Flow cytometric analysis of live cultures after staining for 30min with Hoechst 33342 and thiazole orange allows easy differentiation of uninfected RBCs (red), ring-stage infected cells (blue) and late-stage infected cells (orange) based on RNA and DNA content. Right: 1N ring stages can be used as the reference for calculating genome content of late stages.

**Optional:** At this point, it is possible to cryopreserve infected RBCs for later processing. Harvest cells by centrifugation (800*g* for 3 min at RT) and resuspend in 1 mL of 10 % DMSO prepared in 1× PBS and freeze slowly to −80°C using an isopropanol freezing chamber to minimize cellular damage. Store frozen cells at −80C until you are ready to use the cells, then thaw them quickly by placing cryovials in a 37 °C water bath for 3-5 min. Then transfer the cell suspension to a 1.5 mL microcentrifuge tube and spin at 800*g* for 3 min at RT. Wash twice with 1x PBS before proceeding to step 13.

**CRITICAL:** *From here on, mix gently and avoid vortexing*.

11. Split each sample into 1mL aliquots (one per antibody including isotype controls) in 1.5 mL Lo-bind DNA microcentrifuge tubes and wash twice with 1mL of **Wash buffer** (800 × *g* for 3 min at RT).
12. Resuspend cells in 225 μL **Wash buffer** by gently pipetting.
13. Add 25 μL of activated Concanavalin A-coated magnetic bead slurry to each tube.
14. Incubate at RT on rotator for 10 min at an orbital speed of 10 rpm.

#### Cell permeabilization and antibody binding

**Timing: 2.5 hours**

*Cells attached to conA-beads are permeabilized to allow antibodies to diffuse into the nucleus and bind their targets*.

15. For each antibody make an **Antibody master mix** by combining **Antibody Buffer** with 0.8 μg of antibody to a final volume of 165 μL per sample. For antibodies other than those used in this protocol, final concentrations may have to be determined empirically based on affinity and target abundance.
16. Ensure bead-bound cells are in homogenous suspension by inverting the tube several times, then divide bead-bound cells into aliquots of 250 μL in 1.5 mL microcentrifuge Lo-bind DNA tubes.

**Note**: If antibodies from multiple species are used, include the appropriate isotype control for each species. The amount of antibody needed for optimal signal depends on multiple factors, including its affinity for the target, the abundance of the target and the number of cells used. In our experience high background is more often caused by using too much antibody rather than too little.

17. Place tubes on a magnetic stand to clear then carefully remove and discard supernatant.
18. Remove the tube from the stand and, while holding the tube at an angle, add 150μL of **Antibody master mix** to the side of the tube opposite from the beads. Then tap gently to dislodge the beads.
19. Incubate on a rotator for 2 h at an orbital speed of 10 rpm at RT.

**Pause point**: Antibody incubation may proceed overnight at 4°C.

#### Secondary antibody binding (optional)

**Timing: 1.5 hours**

**Note:** The affinity of proteins A/G to IgG antibodies varies based on host species and IgG subtype (www.bio-rad.com/en-us/applications-technologies/protein-g-affinity). If mouse primary antibodies are being used, it is recommended to use a secondary rabbit-anti-mouse antibody to increase binding to protein A. Although the version of pA/G-MNase sold by Epicypher has an improved antibody compatibility relative to protein A alone, Jenssen and Henikoff suggest that rabbit anti-mouse secondary antibody should still be used in some cases. Otherwise, proceed to step 26.

20. For each mouse antibody make a **Secondary antibody master mix** by combining **Antibody Buffer** with 0.8 μg of Rabbite anti-mouse IgG to a final volume of 165 μL per sample with mouse primary antibody. Remove liquid from cap and side with a quick pulse on a micro-centrifuge, then place on the magnetic stand to clear then remove and discard the supernatant.
21. Add 200 μL of **Dig-wash buffer**, mix by pipetting gently and place on the magnetic stand to clear before removing and discarding the supernatant. Repeat this step two more times
22. Holding the tube on angle and add 150 μL of **Secondary antibody master mix** to the opposite side of the tube from the cells. Tap gently to dislodge the beads.
23. Place the tube on a rotator at 4°C for 1 h at 10 rpm orbital speed.

#### Protein A/G-MNase binding

**Timing: 1 hour**

*Protein A/G-MNase fusion proteins diffuse into the cells and complex with the chromatin-bound antibodies*.

24. Remove liquid from cap and side of the tube with a quick pulse on a micro-centrifuge, then place on the magnetic stand to clear and pull off all the liquid.
25. Add 200 μL of **Dig-wash buffer**, mix by pipetting gently, then place on the magnetic stand to clear and remove all the liquid. Repeat this step two more times.
26. Holding the tube at an angle, add 150μL of the Protein A/G-MNase fusion protein in **Dig-wash buffer** (2.5 μL of 20× stock per sample) directly onto the beads. Tap to dislodge the remaining beads.
27. Place the tube on a rotator at 4°C for 1 h at 10 rpm orbital speed.
28. Place **Incubation buffer** and an aluminum block for 1.5 mL tubes on ice for later use.

#### Chromatin digestion and release

**Timing: 1 hour**

*Chromatin-associated MNase is activated by adding high Ca^2+^* **Incubation buffer**. *Digestion at 0° C prevents the premature diffusion of MNase/DNA complexes and reduces background*.

29. Remove liquid from cap and side with a quick pulse on a micro-centrifuge, then place on the magnetic stand to clear and discard supernatant.
30. Add 200 μL of **Dig-wash buffer**, mix by pipetting gently, then place on the magnetic stand to clear and discard supernatant. Repeat this step two more times.
31. Add 200 μL of **Low-Salt Rinse buffer**, mix by pipetting gently, then place on the magnetic stand to clear and discard supernatant.
32. Add 200 μL of ice-cold **Incubation buffer** directly onto the beads. Tap to dislodge the remaining beads.
33. Incubating at 0°C for 30 min using the pre-chilled aluminum heater block to allow MNase to release bound nucleosomes. Block should be at 0°C to minimize background cleavage.
34. Place on magnetic stand at 4 °C, allow to clear, then remove and *SAVE* the supernatant for possible troubleshooting (see Troubleshooting section at the end)
35. Add 200 μL **Stop buffer** to beads and mix by pipetting gently.
36. Allow liberated nucleosomes to diffuse from the nucleus into the supernatant by incubating for 30 min in a 37°C water bath. **Critical:** *Do not rotate!*
37. Place tube on the magnetic stand to clear. Without disturbing the beads, carefully transfer the supernatant containing liberated chromatin to a fresh Lo-Bind DNA tube.

#### DNA purification by Phenol/Chloroform extraction

**Timing: 2 hours**

*Following the release and diffusion of the nucleosome fragments into the supernatant, DNA is purified and subjected to quality control*.

38. To each sample (200 μL of supernatant containing released CUT&RUN fragments) add 2 μL 10% SDS and 2.5 μL Proteinase K (20 mg/mL). Mix by inversion and incubate 1 h at 50°C in a water bath
39. Add 205μL of phenol-chloroform-isoamyl alcohol 25:24:1. Mix by vortexing at full speed for 5 s.
40. Spin at 16,000*g* for 5 min at RT.
41. Without disturbing the interface, transfer the top liquid phase to a fresh Lo-Bind tube.
42. Add an equivalent volume of chloroform, invert 10-times to mix, then spin at 16,000 *g* for 5 min at RT.
43. Transfer the top liquid phase to a fresh Lo-Bind tube and add 2 μL of 2mg/mL glycogen.
44. Add 500 μL of 100% ethanol and mix 10× by inversion. With an ethanol-resistant pen, mark the bottom of the tube on one side where the pellet is going to form. Chill on ice for 10 min then spin at 16,000 × *g* for 10 min at 4°C with the mark facing outward.
45. Carefully pour off the liquid and dry tube rim by touching it to a paper towel. Rinse the pellet in 1 mL of 100% ethanol and spin at 16,000*g* for 10 min at 4°C.

**Note:** The pellet may not be visible at this point so be careful to not discard the pellet.

46. Carefully pour off the liquid and dry on a paper towel. Let tube air-dry on its side for 5 min.
47. When the pellet is dry, dissolve in 30 μL **TE buffer.**
48. Take 2 μL of each sample to quantify DNA concentration by Qubit Fluorometer using the Qubit dsDNA HS (High Sensitivity) assay kit. In our experience, concentrations at this step are nearly always below 1 ng/μL and have little predictive value of downstream success.
49. Keep the samples on ice at all times.

**Note:** Although samples can be store at −20°C, it is strongly advisable to continue with the protocol until libraries have been PCR amplified.

#### End repair and A-tailing

**Timing: 2 hours**

*The ends of the extracted DNA are repaired and A-tailed to allow subsequent HTS adapter ligation*.

50. Place AMPure XP Beads at RT for later use.
51. In a 1.5 mL microcentrifuge tube prepare the **End repair master mix**.

**End repair master mix**

| Reagent | Amount per reaction* |
| --- | --- |
| EndPrep Enzyme Mix | 1.65 μL |
| EndPrep buffer | 3.85 μL |

\* master mix recipes include 10% extra volume for pipetting error.

52. Combine 5 μL of **End repair master** mix to PCR tubes with 4 ng of DNA template and add ddH2O to a total volume of 30 μL. Mix by vortexing and quick spin to collect at the bottom of the tube.
53. Place the tube in a PCR machine with the heated lid set to ≥ 75° C and run the following program:

| Step | Temperature | Time | Cycles |
| --- | --- | --- | --- |
| End repair | $20^{\circ}\text{C}$ | 30 min | 1 |
| A-tailing | $50^{\circ}\text{C}$ | 60 min | 1 |
| Hold | $4^{\circ}\text{C}$ | forever | |

**Note:** Compared to manufacturer instructions, the A-tailing temperature was decreased to $50^{\circ}\text{C}$ to avoid DNA melting of shorter DNA fragments and the reaction time was increased to 60 min to compensate for lower enzymatic efficiency.

#### Ligation of barcoded HTS adapters

**Timing: 25 minutes**

*Dual barcode HTS adapters are added to the end-repaired and A-tailed fragments*.

54. Remove tube from PCR machine and place on ice.
55. In a 1.5 mL microcentrifuge tube combine the following reagents: **Adapter ligation master mix**

Adapter ligation master mix
| Reagent | Amount per reaction |
| --- | --- |
| NEB ligation Mix | 18 $\mu\text{L}$ |
| NEB ligation Enhancer | 0.6 $\mu\text{L}$ |
| NEXTflex Unique Dual Index Barcode adapters at 0.05 $\mu\text{M}$ | 1.8 $\mu\text{L}$ |

**Note:** We use adapters from NEXTflex Unique Dual Index Barcodes (25 $\mu\text{M}$ concentration) diluted 1:500 in ddH<sub>2</sub>O (0.05 $\mu\text{M}$ final stock concentration). Since adapters are supplied in duplex form avoid heating above RT. Dual index barcodes should be used when sequencing on Illumina systems that use patterned flow cells. If in doubt, use dual index barcodes.

56. Add 17 μL of **Adapter ligation master mix** (final volume of 47 μL) and mix by vortexing. quick spin to collect liquid at the bottom of the tube.
57. Place the tube in the PCR machine with the heated lid OFF and incubate for 15 min at 20° C.

#### Post-ligation Clean-up

**Timing: 20 minutes**

*Excess HTS adapters are removed*.

58. Transfer the samples into a 1.5 mL Lo-Bind microcentrifuge tube.
59. Add 80μL of AMPure XP bead slurry (1.7× ratio). Mix well by pipetting at least 10 times and incubate for 5 min at RT.

**Note:** Prior to use equilibrate AMPure XP beads to RT for at least 30 minutes.

60. Place the tube on the magnetic stand to separate beads from liquid. Discard the supernatant.
61. Wash twice with 200 μL 80% Ethanol while keeping the tubes on the magnetic stand.
62. Discard supernatant and let dry for up to 5 min

**Critical:** *DO NOT DISCARD THE BEADS! Tubes must remain in the magnetic stand the whole time. Avoid overdrying the beads. Add **TE buffer** to the beads when all visible liquid has evaporated but beads are still dark brown and glossy looking*.

63. Resuspend beads in 15 μL of **TE buffer**. Mix by pipetting 10 times and incubate for 5 min at RT.
64. Place the tube back on the magnetic stand. Allow to clear and transfer the supernatant (^~^13 μL) to a fresh PCR tube.

#### Library PCR amplification

**Timing: 35 minutes**

*Fragments are PCR amplified using KAPA polymerase, which is more efficient at amplifying AT-rich fragments*.

65. In a 1.5 mL microcentrifuge tube combine the following reagents:

PCR master mix
| Reagent | Amount per reaction |
| --- | --- |
| 2x KAPA HotStart ready mix | 27.5 µL |
| NEXTflex Primer mix | 5.5 µL |
| ddH<sub>2</sub>O | 7.7 µL |

**Note:** We use NEXTflex Primer Mix (12.5 μM stock concentration) that comes with NEXTFLEX Unique Dual Index barcodes sets, at a final concentration of 1.2 μM.

66. Add 37 μL of **PCR master mix** to each sample for a final volume of 50 μL. Mix by vortexing and then quick spin to collect liquid at the bottom of the tube.
67. Place in a PCR machine with the heated lid set to 100°C and run the following program: **PCR cycling conditions**

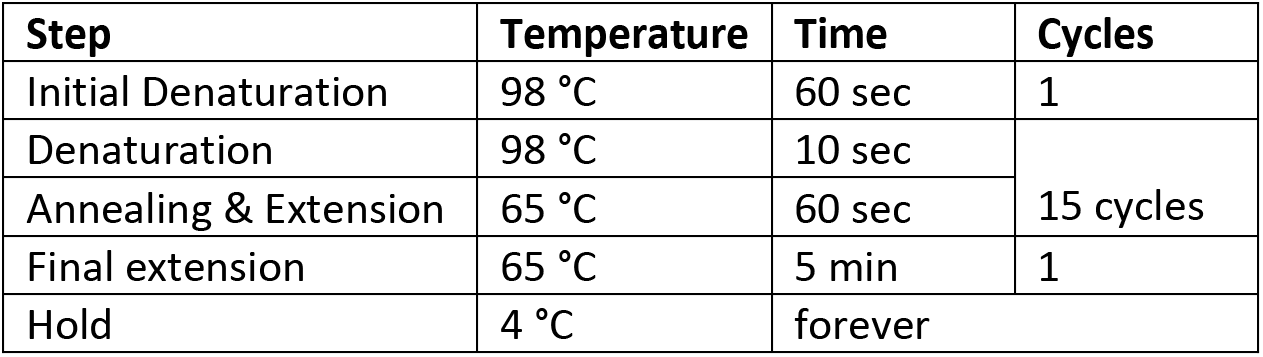

**Pause point**: Samples can be stored at −20°C until further use.

#### Clean-up of PCR amplified library

**Timing: 35 minutes**

*Excess primers are removed*.

**Note:** This round of selection is aimed to remove primer and adapter pairs below 150 bp.

68. Add 50μL of AMPure XP bead slurry (1.0× ratio) to each amplified library and mix by pipetting at least 10 times, then incubate for 5 min at RT.
69. Place the tube on the magnetic stand for 5 min to separate beads from supernatant.
70. After the solution is clear, carefully remove and discard the supernatant. Be careful not to disturb the beads that contain DNA targets.

**Critical:** *DO NOT DISCARD THE BEADS! Tubes must remain in the magnetic stand the whole time*.

71. Add 200 μL of 80% Ethanol (freshly prepared) to the tube while in the magnetic stand. Incubate at RT for 30 seconds, and carefully discard the supernatant. Repeat this step for a total of two washes and remove all visible liquid after the second wash.
72. Air dry the beads for up to 5 min while the tube is on the magnetic stand with the lid open. *Avoid over-drying the beads*.
73. Remove the tube from magnetic stand. Elute the DNA fragment from the beads by adding 15 μL of **TE buffer**. Mix well by pipetting up and down 10 times. Incubate for 2 min or more at RT.
74. Place the tube on the magnetic stand. After 5 min (or when the solution is clear) transfer the supernatant (^~^13 μL) to a new fresh microcentrifuge Lo-Bind tube.

**Pause point**: Samples can be stored at −20° C until further use.

#### Library quality control and Illumina sequencing

**Timing: 25 min**

75. Take 2 μL of each sample to quantify DNA concentration by using fluorescence detection Qubit Fluorometer using the Qubit dsDNA HS (High Sensitivity) assay kit. For the antibodies/samples in this example, concentrations were usually in the 2-15 ng/μL range (lower for Isotype controls).
76. Use 1 ng of each library to check fragments size distribution and molar concentration using a Bioanalyzer or TapeStation system. (See Figure 7).

**Figure 7.**
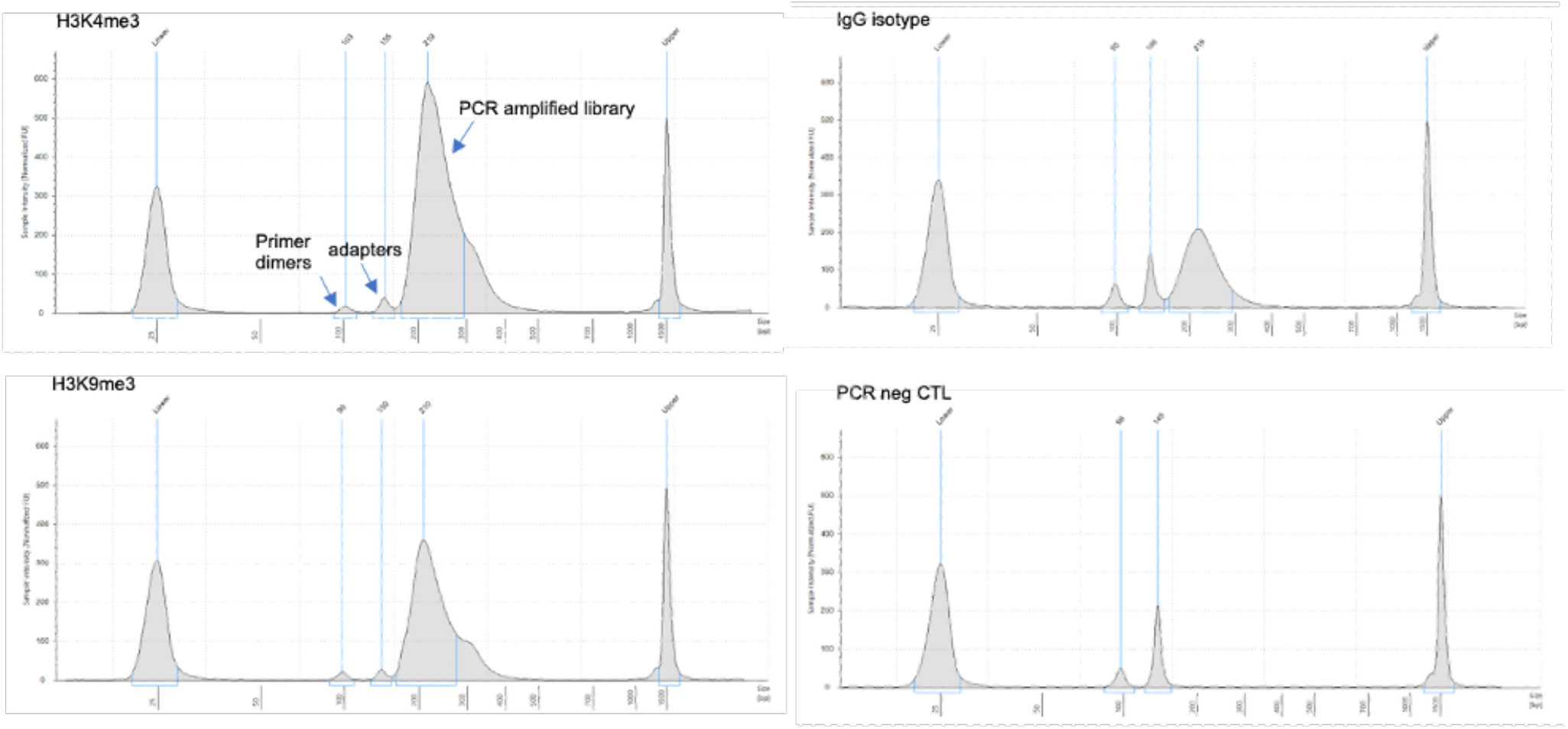
Example size distributions of CUT&RUN libraries from H3K9me3, H3K4me3, IgG isotype, and PCR negative control as analyzed by TapeStation. Primer dimer, adapter, and library peaks are indicated in the top left panel.

**Note:** If the fraction of adapters dimers exceeds 20% of your library, an additional round of size selection is required. Do NOT pool libraries with high adapter content with high quality libraries as this will require size selecting the pool and reduce the abundance of high-quality libraries.

77. Pool libraries at desired ratio reflecting the desired relative sequencing depths for each library (In this case we combined equimolar amounts (0.02 pmol) of 25 libraries to generate the library pool).
78. Rerun pooled libraries in a Bioanalyzer or TapeStation to check size profile and molar concentration.
79. Perform paired-end Illumina sequencing following manufacturer’s instructions. 50 bp paired-end reads are sufficient for accurate mapping.

## Expected outcomes

When starting with 1×10^7^ schizont stages with an average nuclear content of 3-5N, post-clean-up library yield was in the range of 10-40 ng after 15 cycles of PCR. Figure 2 shows examples of CUT&RUN library size profiles obtained with anti-H3K9me3 and anti-H3K4me3 antibodies. The ratio of primer dimers and adapters in our libraries was around 4%, which is 5x lower than normally reported for these assays. Library size profile obtained with rabbit IgG antibody represents unspecific bound regions that have been PCR amplified. Also, a PCR negative control with no template DNA was shown, to dismiss this sort of contamination during library preparation.

## Quantification and statistical analysis

The libraries shown in these examples were pooled with other CUT&RUN libraries at equimolar ratios and sequenced using an Illumina NextSeq2000 sequencer in paired-end 50bp P2 mode. We initially aimed at a target of 10 million PE50 read pairs per library to ensure good coverage. Subsequent down-sampling analysis showed that as low as 1 million read pairs provides excellent signal-to-noise ratios for these antibodies (Figure 4).

For the comparison of the CUT&RUN profiles to published profiles (Figure 3) we obtained raw read files from public repositories using SRAtools (https://github.com/ncbi/sra-tools). H3K9me3 and input raw ChIP-seq reads from (Bunnik et al. 2018) were downloaded from NCBI Sequence Read Archive (accession number SRP091939). *Pf*HP1 and input raw ChIP-seq reads from (Fraschka et al. 2018) were downloaded from NCBI’s Gene Expression Omnibus (accession number GSE102695). H3K4me3 and input raw ChIP-seq reads from (Bartfai et al. 2010) were downloaded from NCBI’s Gene Expression Omnibus (accession number <u>GSE210062</u>).

Raw reads were quality trimmed using Trimmomatic v0.38 (Bolger et al. 2014) to remove residual adapter sequences and low quality leading and trailing bases to retain properly paired reads of length >=30 bases after trimming, as previously published (Campelo Morillo et al. 2022). Trimmed reads were aligned to PlasmoDB version 51 *P. falciparum* 3D7 reference genome (Warrenfeltz et al. 2018) using BWA v.0.7.1 (Li & Durbin 2009). SAMtools v.1.10 (Li et al. 2009) was used to sort and index BAM files. Normalized fold enrichment tracks were generated by using the MACS2 v.2.2.7.1 (Zhang et al. 2008) callpeak function with settings: -f BAMPE -B -g 2.3e7 -q 0.05 –nomodel –broad –keep-dup auto –max gap 500. Bedgraph outputs were then passed into the bdgcmp function with the setting -m FE (fold enrichment) to generate signal tracks to profile histone modification enrichment levels compared to either input (for ChIPseq) or isotype IgG control (CUT&RUN). Using the bedGraphToBigWig tool (Kent et al. 2010), we converted the genome-wide coverage and enrichment tracks from bedGraph into BigWig format. These files are available for download at NCBI Gene Expression Omnibus (GEO) under accession number <u>GSE210062</u> and can be directly loaded into the PlasmoDB genome browser (Warrenfeltz et al. 2018) for detailed inspection.

Coverage and enrichment tracks were visualized using Integrative Genome Viewer IGV (Thorvaldsdóttir et al. 2013) or the GenomicRanges v.1.44.0 and GViz Viz v1.38.1 (Hahne & Ivanek 2016) R packages within the Bioconductor project (release 3.13) (Huber et al. 2015). The full analysis pipeline can be found at https://github.com/KafsackLab/PfCUTandRUN. Representative tracks of H3K9me3, H3K4me3, and HP1 produced using CUT&RUN or ChIP-seq at chromosome 8 of *P. falciparum* are shown in Figure 4.

## Troubleshooting

### Problem 1: Low DNA concentration following library amplification.

If DNA is low or undetectable in step 84, check the following:

-beads were not activated properly
-beads got stuck on tube lid or walls during incubation steps
-diffusion of MNase/DNA/antibody complex occurred prematurely

#### Potential solution

Remember to activate beads before initiating the protocol. Give a quick spin after each incubation to recover beads from tube lid and walls. Try not to pipette too harshly since cells can be very fragile after permeabilization. Always perform the digestion at 0°C and do not agitate the tube once High Ca^2+^ **Incubation buffer** is added to the isolated DNA from supernatants saved in step 36. No DNA should be detected in this fraction, if so, this is indicative of premature diffusion of nucleosomes during digestion. For low abundance targets, increasing the cycle number during PCR library amplification may help.

### Limitations

This protocol relies on the availability of a specific antibody against histones PTM that would be able to recognize correspondent modification in *Plasmodium falciparum*. This should be tested empirically by Western blot or immunoprecipitation assays. We have only used rabbit antibodies. Antibodies generated in other species should be tested according to experimental set up and availability and antibodies raised in other species may benefit from the inclusion of rabbit secondary antibodies. This protocol was optimized for use with abundant chromatin marks on nucleosomes. For rare modifications, or non-nucleosomal targets, like transcription factors and histone modifiers, further optimization may be required.

## Resource availability

### Lead contact

Further information and requests for resources and reagents should be directed to and will be fulfilled by the lead contact, Björn F.C. Kafsack.

### Materials availability

No new materials were generated in this study.

### Data and code availability

The full analysis pipeline can be found at https://github.com/KafsackLab/PfCUTandRUN.

Raw (fastq) and processed (BigWig) high-throughput sequencing data have been deposited in the NCBI Gene Expression Omnibus under accession number <u>GSE210062</u>.

## Acknowledgments

We thank the VEuPathDB team and the Weill Cornell Medicine genomics core for technical support. This work was supported by funding from Weill Cornell Medicine, NIH NIAID 1R01 AI141965 and NIH NIAID 1R01 AI138499, Alice Bohmfalk Charitable Trust, and Charles A. Frueauff Foundation.

## Author contributions

Conceptualization by B.F.C.K. and R.A.C.M.; R.A.C.M. and C.T.H. carried out investigation; Software, formal analysis and data curation were provided by B.F.C.K. and C.T.H.; R.A.C.M. wrote the original draft with contributions from C.T.H. and K.K.; B.F.C.K. and R.A.C.M. reviewed and edited the article; Visualization was conducted by R.A.C.M., C.T.H. and B.F.C.K.; Project administration and funding acquisition were carried out by B.F.C.K.

---

**Supplementary Figure 1.**
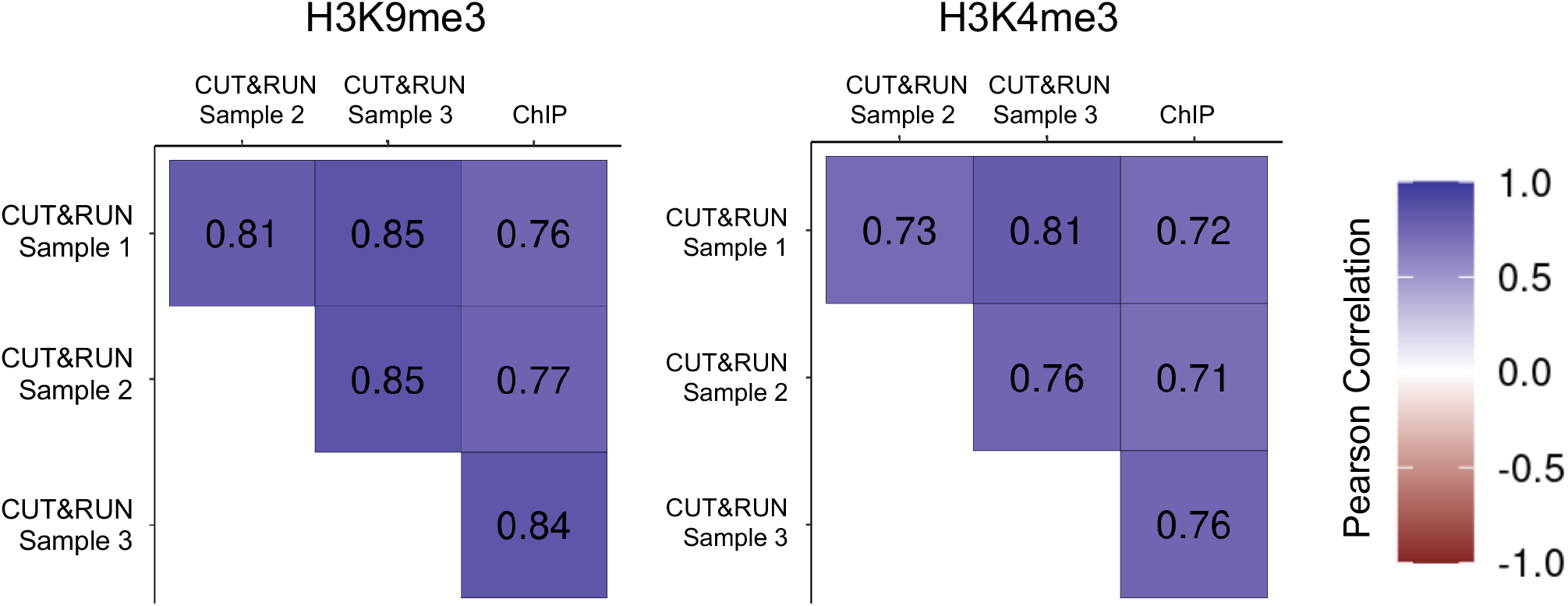
Correlation of fold-enrichment between CUT&RUN replicates and published ChIP-seq data. Color and values shown indicate the genome-wide Pearson’s correlation coefficient.

**Supplementary Figure 2.**
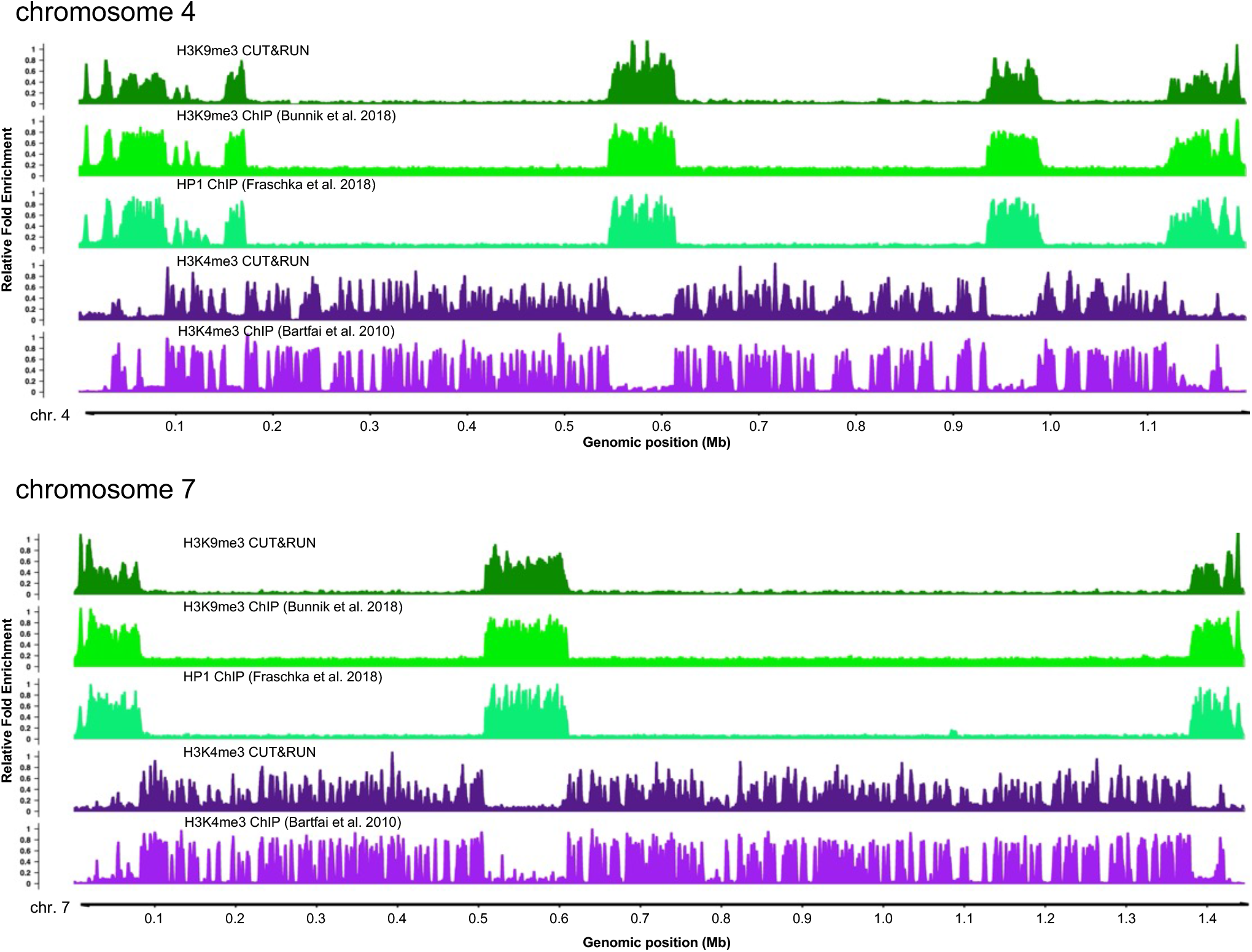
Comparison of H3K4me3 and H3K9me3 profiles obtained by CUT&RUN to published ChIP-seq profiles for full length chromosomes 4 and 7. Tracks show the relative fold-enrichment of specific histone modifications, H3K9me3 & H3K4me3, as well as for heterochromatin protein 1, the H3K9me3 histone reader, versus either isotype controls for CUT&RUN or input DNA (ChIPseq). Sequence reads from published stage-matched H3K9me3 (SRR4444647, SRR4444639), HP1 (SRR5935737, SRR5935738), and H3K4me3 (SRR065659, SRR065664) ChIP-seq experiments were downloaded from the NCBI Sequence Read Archive.

